# The yolk sac vasculature in early avian embryo provides a novel model for the analysis of cancer extravasation

**DOI:** 10.1101/2024.05.12.593798

**Authors:** Mizuki Morita, Ryo Fujii, Asuka Ryuno, Manami Morimoto, Akihito Inoko, Takahiro Inoue, Junichi Ikenouchi, Yuji Atsuta, Yoshiki Hayashi, Takayuki Teramoto, Daisuke Saito

## Abstract

Hematogenous metastasis, a key trait of cancer cells, involves a complex sequence of cell migration steps, including intravasation, circulation, arrest in capillary vessels, and extravasation. Among these steps, extravasation is challenging to image in amniotes like humans and mice due to its unpredictable timing and location, limiting our understanding of cellular and molecular mechanisms through imaging. Establishing a new cancer carrier model with high-resolution imaging capabilities in amniotes is crucial. In this study, we investigated the yolk sac vasculature (YSV) of early avian embryos (chickens and quail) as a new model for studying extravasation, offering excellent imaging capabilities. We examined the YSV structure and attempted fluorescent labeling to enhance visibility. We then injected mCherry-labeled HT-1080 cells into YSV and observed their behavior, revealing distinct morphologies and extravasation dynamics. Our findings suggest that the YSV model holds promise as a novel cancer carrier model for elucidating cellular and molecular mechanisms through imaging-based approaches.

## INTRODUCTION

Metastasis stands a pivotal event in cancer progression, wherein malignant cells disseminate from the primary tumor to distant sites, primarily via the bloodstream. This intricate process unfolds in five distinct stages: (1) epithelial-mesenchymal transition (EMT) fostering enhanced cellular motility; (2) cancer cell translocation into the intravascular space, known as intravasation; (3) circulation through the bloodstream; (4) arrest of cancer cells at the capillaries; and (5) trans-endothelial migration (TEM) into the surrounding tissue (Katt et al., 2018; Reymond et al., 2013; Sahai, 2007; Steeg, 2016). The latter two steps collectively constitute extravasation.

While considerable progress has been made in targeting steps 1-3 (Giampieri et al., 2009; Gregory et al., 2008; Minder et al., 2015; Yankaskas et al., 2021; Yuan et al., 2014), our understanding and therapeutic strategies for extravasation remain constrained. Its occurrence is sporadic in both living humans and murine mice, rendering prediction challenging. Additionally, the constant physiological movements, such as respiration, cardiac pulsations, and voluntary motions, complicate live imaging-based analyses. Consequently, alternative analytical approaches have emerged.

Microfluidic device models, integrating vascularized endothelial cells and fluid flow, have illuminated the roles of selectins and integrins in the arrest phase, along with the contributions of accompanying cells, fluid dynamics, and TNF stimulation in TEM (Burdick et al., 2006; Chen et al., 2016b; Shin et al., 2011). Zebrafish embryonic vasculature models have delineated a cancer-driven TEM facilitated by invadopodia (Stoletov et al., 2007), alongside an endothelium-driven TEM termed angiopellosis, where vascular endothelial cells engulf cancer cells (Allen et al., 2017; Follain et al., 2018). The avian embryo chorioallantoic membrane (CAM) model has similarly demonstrated invadopodia-mediated TEM, with Tks and MMP dependency (Leong et al., 2014).

Despite the advantages of these imaging models, such as simplicity, transparency in living organisms, and membership within the amniotic class akin to mammals, they have limitations including *in vitro* settings, reduced body temperatures, and relatively lower imaging resolutions. Thus, mutual validation among these models is essential. The emergence of amniotic organism models facilitating high-quality live imaging holds promise for significant utility in this domain.

In this study, we aimed to validate whether the yolk sac vasculature (YSV) of avian embryos could serve as a model meeting these criteria. The YSV is an extraembryonic vascular system that surrounds the early avian embryos. It comprises a small, closed vascular network with a diameter of approximately 30 mm, unfolding in a two-dimensional plane with a vein or capillary-like structure. Due to its small and flat structure, coupled with the advantage of *ex vivo* whole culture (Chapman et al., 2001; Kidokoro et al., 2018; New, 1955), it enables capturing nearly all positions of transplanted cancer cells while capturing individual cellular behaviors during extravasation at high resolution. In this study, we successfully captured several cellular behaviors exhibited by cancer cells during extravasation. Additionally, we demonstrated the effectiveness of this system as a quantitative evaluation tool for cancer extravasation. Our finding leads us to conclude that YSV is a valuable new model for extravasation analysis.

## MATERIALS and METHODS

### Animals, staging, and animal care

Fertilized chicken (*Gallus gallus domesticus*, White leghorn) eggs and fertilized quail (*Coturnix japonica*) eggs were obtained from Takeuchi poultry farm (Nara, Japan) and from Nagoya University through the National Bio-Resource Project of the MEXT, Japan, respectively. Eggs were incubated at 38.5℃, and embryos were staged according to Hamburger and Hamilton’s criteria (Hamburger and Hamilton, 1951) or Eyal-Giladi and Kochav’s criteria (Eyal-Giladi and Kochav, 1976). All animal experiments were conducted with the approval of the Institutional Animal Care and Use Committees at Kyushu University.

### Plasmid constructions

pT2A-BI-TRE-ZsGreen1: The full-length of ZsGreen1 was amplified by PCR, and subcloned into the EcoRI-PstI site of pT2A-BI-TRE (Saito et al., 2022). pT2A-BI-TRE-mCherry: The full-length of mCherry was amplified by PCR, and subcloned into the EcoRI-BglII site of pT2A-BI-TRE. pT2A-BI-TRE-LAP2bcGFP-Lifeact-mCherry: Lifeact-mCherry was isolated from pT2A-CAGGS-Lifeact-mCherry vector digested by MluI-NheI and subcloned into pT2A-BI-TRE which had been treated either with MluI-NheI. The fragment of rat LAP2b-AcGFP was amplified from pAcGFP-N1 (Addgene Plasmid #62044) by PCR and subcloned into the EcoRI-BglII site of pT2A-BI-TRE-Lifeact-mCherry. pT2A-BI-TRE-ZsGreen1-ΔC-Integrinβ1: The ΔC-Integrinβ1 (Yoshino et al., 2014) was subcloned into the MluI-EcoRV site of pT2A-BItight-ZsGreen1-TRE.

### Cells culture

HT-1080 cells (JCRB Cell Bank) were maintained in Dulbecco’s modified Eagle’s medium (DMEM; Nakarai tesque) containing 4.5g/l Glucose with L-Gln and Sodium Pyruvate supplemented with 10% (v/v) fetal bovine serum (FBS; Gibco), Gluta-Max^TM^-1 (Gibco), and 1% (v/v) penicillin-streptomycin-ampphotericin (PSA; FUJIFILM Wako Pure Chemical Corporation) at 37°C and 5% CO_2_. Cells were passaged using 0.05% trypsin-EDTA (Gibco).

### Plasmid transfection and establishment of gene-manipulated cells

1 × 10^6^ cultured cancer cells were washed with OPTI-MEM (Gibco) and seeded in a 6-well plate containing 1 ml of medium without PSA for 18-24 h (overnight). A transfection mix containing 4.0 μg of total plasmids and 7.0 μl of Lipofectamine 2000 (ThermoFisher Scientific) in 250 μl of OPTI-MEM was added to cancer cells in the 6-well plate. We utilized 4 different sets of plasmids for transfection to cancer cells: pT2A-BI-TRE-ZsGreen1 + pT2A-CAGGS-Tet3G-2A-PuroR (Saito et al., 2022) + pCAGGS-T2TP (Saito et al., 2022); pT2A-BI-TRE-LAP2bcGFP-Lifeact-mCherry + pT2A-CAGGS-Tet3G-2A-PuroR + pCAGGS-T2TP; pT2A-BI-TRE-mCherry + pT2A-CAGGS-Tet3G-2A-PuroR + pCAGGS-T2TP; pT2A-BI-TRE-ZsGreen1-ΔC-Integrinβ1 (Itgnβ1) + pT2A-CAGGS-Tet3G-2A-PuroR + pCAGGS-T2TP. The medium was replaced with conventional medium containing antibiotics 6 hours after addition of the transfection mix. After culturing with medium containing antibiotics for 2 days, the cancer cells were further cultured in the medium containing 2-5 μg/ml puromycin for 3 days to enrich puromycin-resistant cells. Following puromycin treatment, ZsGreen1-expressing cells were sorted by an SH800 cell sorter (Sony) based on the positivity of ZsGreen1 signals.

### Vasculature labeling using dextran, acetylated low-density lipoprotein, and Lens culinaris lectin

For fluorescence staining of the vasculature, either 2% fluorescein isothiocyanate-dextran (FITC-Dex) (average mol wt 250,000) (Sigma-Aldrich, FD250S) in OPTI-MEM, or 50% Alexa Fluor^TM^ 594 acetylated low-density lipoprotein (594-AcLDL) (ThermoFisher Scientific, L35353) in OPTI-MEM, or 20% Rhodamine-Lens culinaris lectin (Rho-LCA) (Vector Laboratorries, L-1040) in OPTI-MEM, was injected into the embryo’s heart using a fine glass capillary before live-imaging or fixation.

### Cancer cell injection into embryos and *ex vivo* live-imaging

Cancer cells on the culture dish were detached using 0.05% trypsin-EDTA/PBS solution. The trypsin reaction was halted by adding an equal volume of fetal bovine serum (FBS) to the trypsin solution. After centrifugation at 1,500 rpm for 5 minutes, the cells were washed several times with OPTI-MEM. The number of viable cells was adjusted to 20,000 cells/μl for each cell type, mixed equally, and 1 μl of this suspension was injected into the hearts of HH15 quail embryos using a fine glass capillary. For tet-on induction, 1 μg/ml Doxycycline (Dox) (Clontech) was added to the cultured cells’ medium 24 hours before injection. The embryos were then incubated at 38.5°C until the intended developmental stage. For live imaging, manipulated embryos were delicately removed from the eggs using filter paper rings and positioned dorsal-side down on 50 mm glass bottom dishes pre-coated with thin albumen (Chapman et al., 2001; Kidokoro et al., 2018). The embryos were cultured at 38.5°C for further observation.

### Fixation of embryos and immunofluorescence staining

Whole embryos were fixed in 4% PFA/PBS (paraformaldehyde/phosphate buffered saline) at 4 ℃ overnight. For immunostainings involving QH-1, alpha smooth muscle actin (αSMA), and laminin in quail whole YSV, samples underwent the following procedure: Washed six times for 30 minutes each with TNTT (0.1 M Tris-HCl (pH 7.5), 0.15 M NaCl, 0.05% Tween 20, 0.1% TritonX-100) at 4℃. Immersed in a 3% hydrogen peroxide methanol solution at 4 ℃ for 2days to inactivate endogenous peroxidase. Washed three times for 30 minutes each in TNTT at 4 ℃ to remove methanol.Blocked with 1% blocking reagents (Roche) in TNTT buffer for 2 hours at 4 ℃ to prevent no specific binding. Incubated overnight at 4 ℃ with the primary antibodies diluted in 1% blocking solution: (1) anti-αSMA rabbit polyclonal antibody (GeneTex, GTX100034 1:500) and QH1 antibody (Hybridoma Bank, AB_531829, 1:200),or (2) anti-laminin rabbit polyclonal antibody (Abcam, ab11575, 1:250) and QH1 antibody (Hybridoma Bank, AB_531829, 1:200). Washed three times for 30 minutes each in TNTT at 4 ℃ to remove unbound primary antibodies. Incubated overnight at 4 ℃ overnight with secondary antibodies diluted in 1% blocking solution: RT for 1 hour; (1) 1:500 anti-Mouse IgG-HRP-linked whole ab sheep antibody (Cytiva NA931V) and 1:500 anti-rabbit IgG-Alexa 488-conjugated donkey antibody (Invitrogen A21206), (2) 1:500 anti-Mouse IgG-HRP-linked whole ab sheep antibody (Cytiva NA931V) and 1:500 anti-rabbit IgG-Alexa 647-conjugated donkey antibody (Invitrogen A31573). Washed three times for 30 minutes each in TNTT at 4 ℃ to remove secondary antibodies. Carried out fluorescence sensitization by incubating the samples in 1% Cyanine 5 regent (Perkin Elmer FP1171) with amplification diluent (Perkin Elmer FP1135) for 15 minutes at room temperature (RT). Rendered embryos transparent, if necessary, by immersing the sensitized YSV in 40% glycerol/PBS for 1 hour at 4 ℃, followed by treatment with 60% glycerol/PBS for 2-3 hours at 4 ℃, and finally placed in 80% glycerol /DDW at 4℃ overnight.

For immunostainings involving QH1, ZsGreen1, and mCherry in quail whole YSV, samples underwent the following procedure: Washed six times for 30 minutes each with TNTT (0.1 M Tris-HCl (pH 7.5), 0.15 M NaCl, 0.05% Tween 20, 0.1% TritonX-100) at 4℃. Immersed in a 3% hydrogen peroxide methanol solution at 4 ℃ for 2days to inactivate endogenous peroxidase. Washed three times for 30 minutes each in TNTT at 4 ℃ to remove methanol. Blocked with 1% blocking reagents (Roche) in TNTT buffer for 2 hours at 4 ℃ to prevent no specific binding. Incubated overnight at 4 ℃ with the primary antibodies diluted in 1% blocking solution: QH1 antibody (DSHB, AB_531829, 1:200), anti-GFP goat polyclonal antibody (St John’s Laboratory, STJ140005, 1:400), and anti-RFP rabbit polyclonal antibody (ROCKLAND, 600-401-379, 1:500). Washed three times for 30 minutes each in TNTT at 4 ℃ to remove unbound primary antibodies. Incubated overnight at 4 ℃ overnight with secondary antibodies diluted in 1% blocking solution: 1:500 anti-Mouse IgG-HRP-linked whole ab sheep antibody (Cytiva NA931V), 1:500 anti-rabbit IgG-Alexa 555-conjugated donkey antibody (Invitrogen A31572), and 1:500 anti-goat IgG-Alexa 488-conjugated donkey antibody (Jackson Immuno Research 705-545-147). Washed three times for 30 minutes each in TNTT at 4 ℃ to remove secondary antibodies. Carried out fluorescence sensitization by incubating the samples in 1% Cyanine 5 regent (Perkin Elmer FP1171) with amplification diluent (Perkin Elmer FP1135) for 15 minutes at room temperature (RT). Rendered embryos transparent, if necessary, by immersing the sensitized YSV in 40% glycerol/PBS for 1 hour at 4 ℃, followed by treatment with 60% glycerol/PBS for 2-3 hours at 4 ℃, and finally placed in 80% glycerol /DDW at 4℃ overnight.

### Image acquisition and processing

*In ovo* live imaging of cultured embryos was performed using the MVX10 microscope (Olympus) or the Leica S9 D Stereo Microscope (Leica) and acquired using high speed recording software (Hamamatsu) or Flexacam C 3 (Leica). The acquired images were enlarged, edited using Fiji-ImageJ (NIH).

For *ex vivo* live imaging of cultured embryos, the samples on the glass bottom dishes (Matsunami) were placed in the incubation chamber (37°C, 5% CO_2_) (TOKAIHIT STX) equipped with the inverted microscope, IX83 (Olympus) interfaced to a spinning-disk confocal microscopy (Dragonfly200, OXFORD Instruments). Images were captured with a device camera and acquired using Fusion software (Dragonfly200, OXFORD Instruments). The acquired images were enlarged, edited using the built-in function of Imaris (OXFORD Instruments) or Fiji-ImageJ (NIH).

Image acquisition of fixed whole-mount images was obtained with the confocal laser scanning microscope (Leica TCS SP8, Leica Microsystems) or IX83 (Olympus) interfaced to a spinning-disk confocal microscopy (Dragonfly200, OXFORD Instruments). Images were captured with a device camera and acquired using LAS X Life Science Microscope Software Platform (Leica Application Suite X (LAS X), Leica) or Fusion software (Dragonfly200, OXFORD Instruments). Acquired Z-series images were processed for 3D reconstruction using Imaris software (ver.9.5, Oxford Instruments). These images were further reconstructed into 3D iso-surfaces with texture, and clipped with appropriate dorsal-ventral plane(s).

### Quantification and statistical analysis

Box plots represent the mean, upper and lower interquartile, error bars (s.e.m) with median (x). p values were obtained by a 2-tailed, unpaired Student’s t test (Excel). All bar graphs and line graphs represent the mean, standard error. p values were obtained by a 2-tailed, unpaired Student’s t test (Excel).

## RESULTS

### YSV characterization

In evaluating the YSV’s potential as a model for analyzing cancer extravasation, we initially scrutinized its characteristics. The YSV comprises extraembryonic vessels that radiate concentrically around the embryo proper, encompassing all vessels in chick and quail embryos at Hamburger and Hamilton’s stage (HH) 15 within an area of approximately 15 mm^2^ radius (Fig. 1A). Among these vessels, the vitelline artery (VA) stands out as the thickest, connecting to the dorsal aorta at the lateral level within the embryo proper and facilitating the expulsion of blood from the heart into the entire YSV. Gradually tapering towards the periphery, the VA displays numerous branching points. Blood returns to the heart via either the sinus vein (SV) or anterior vessels (Fig. 1B, C).

**Figure 1.**
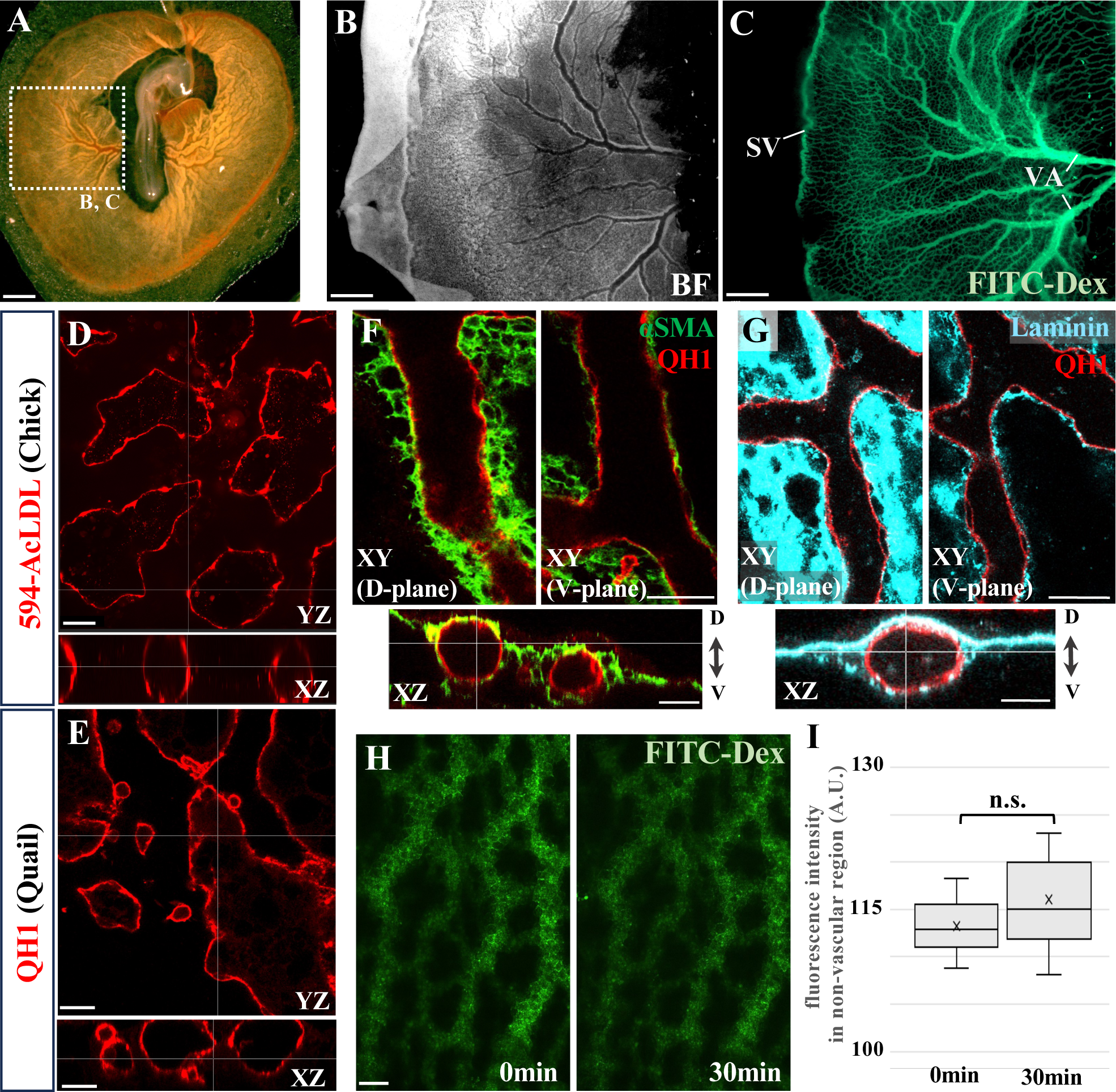
YSV characterization. **(A-C**) Dorsal view of HH15 chick embryo injected with FITC-Dex. (A) is whole image (bright field), (B) (bright field) and (C) (FITC-Dex) are corresponded to lateral YSV region, magnified views of dotted box area shown in (A). SV, sinus vein; VA, vitelline artery. **(D, E)** Vascular patterns in lateral YSV regions in HH15 chick injected with 594-AcLDL (D) and HH15 quail stained with QH1 antibody (E). XYZ means axis. **(F, G) A**lpha-smooth muscle actin (αSMA: green), Laminin (cyan), and QH-1 (red) immunofluorescence pictures in lateral YSV regions of HH15 quail embryos. D, dorsal; V, ventral. **(H, I)** Images (H) and quantification (I) of fluorescence intensity of FITC-Dex in non-vascular region of HH16 chick embryo (N=3) after FITC-Dex injection. Scale bars: 1000 μm in A, B, C, 40 μm in **D, E, F, G** 50 µm in **H**.

Given its small size and flat nature, confocal imaging of the YSV and its three-dimensional reconstruction can be achieved through whole-mount endothelial staining. In both chick and quail embryos at HH15, individual blood vessels in this lateral region of the YSV, particularly conducive to observation, indicate an average diameter of approximately 15-100 μm (Fig. 1D, E), maintained from HH15 to HH16.

Structurally, at HH15, the YSV exhibits a typical vascular arrangement with endothelial cells (ECs) forming the innermost layer, surrounded by basement membrane (BM) and smooth muscle cells (SMCs) (Fig. 1F, G). Notably, the vascular wall facing the dorsal (albumen) side displays a thick, continuous BM and SMC layer, whereas the ventral (yolk) side shows less BM and SMC presence (Fig. 1 F, G). This vascular architecture remains consistent through development to HH16. Tracking analysis of infused microbeads indicates that blood flow velocity varies depending on location, ranging from approximately a minimum of 40 μm/s to a maximum of 350 μm/s (data not shown). Analysis of vascular leakiness using 250,000 MW fluorescein isothiocyanate-dextran (FITC-Dex) revealed minimal leakage even after 30 min post-administration (Fig. 1H, I).

Based on these findings, it was concluded that the YSV resembles veins or capillaries, exhibiting a degree of vascular integrity and distinct characteristics along the dorsal-ventral axis. Given that cancer cell extravasation often occurs in capillaries or veins (Kalluri, 2003; Li et al., 2020), the YSV appears promising as a model for analyzing cancer cell extravasation.

### Vasculature visualization in living and fixed chick and quail embryo

To elucidate cancer cell extravasation, visualizing the YSV in living embryos at the HH15-16 stages is essential. We meticulously assessed three methodologies for this purpose. Firstly, we implemented the injection of FITC-Dex into vasculature, following the protocol by Entenberg *et al*. (Entenberg et al., 2017) (Fig. 2A, B), as also depicted in Fig.1. This approach enables the observation of blood flow dynamics and distinctive demarcation between blood plasma and endothelium (Fig. 2B, D, E). However, direct visualization of ECs is not achieved, and fading occurs rapidly, disappearing within approximately 30 minutes under fluorescence observation (Fig. 2G, I). Secondly, we employed the administration of Alexa Fluor^TM^ 594-conjugated AcLDL (594-AcLDL) in the vasculature, as per the method described by Lu *et al*. (Lu et al., 2007) (Fig. 2A, C). This results in the uptake of 594-AcLDL by ECs, manifesting as dot-like signals within these cells (Fig. 2D, F). This facilitates visualization of the approximate contour of ECs, which can be sustained for at least 6 hours (Fig. 2H, I). Thirdly, we verified the effectiveness of YSV visalization through the injection of Rhodamine-Lens culinaris lectin (Rho-LCA) into vasculature, following protocol by Leong *et al*. (Leong et al., 2014). Rho-LCA injection allows clear visulalization of endothelial cells (Fig. 2J, K), but blood stasis occurs due to erythrocyte aggregation (Fig. 2L). Injection at concentrations that do not cause erythrocyte aggregation fails to adequately label ECs. In conclusion, vascular labeling methods by FITC-Dex and 594-AcLDL can be utilize depending on the specific application, and they can be used simultaneously in living chick and (or) quail embryos without erythrocyte aggregation and hemodynamics stasis, but Rho-LCA is not suitable for live analysis at HH15-16.

**Figure 2.**
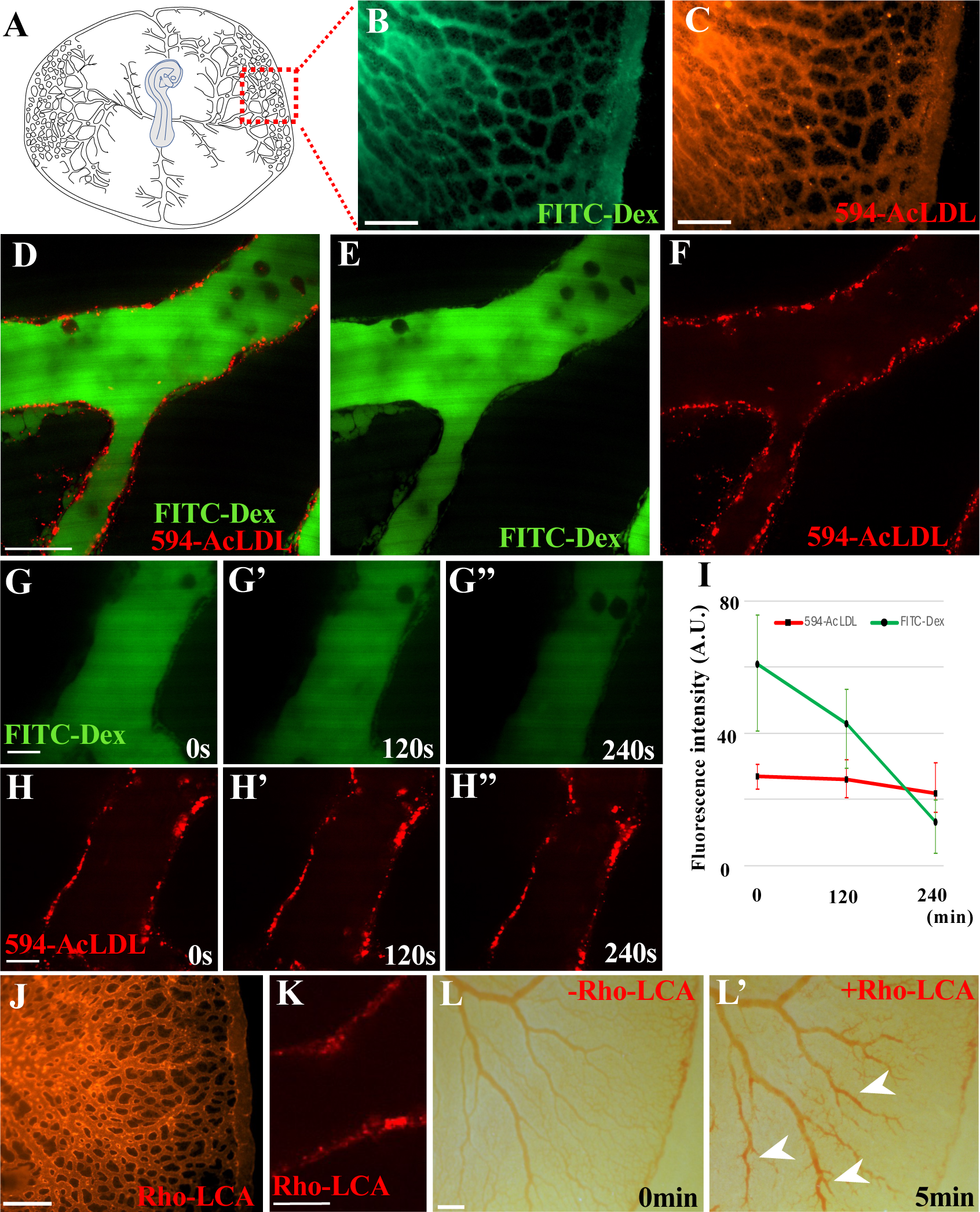
Vasculature visualization in living chick and quail embryo. **(A)** Dorsal view of HH15 avian embryo. **(B, C)** Images in (B) (FITC-Dex) and (C) (594-AcLDL) are corresponded to red dotted box area shown in (A) after 15 minutes post-injection. **(D-F)** The magnified views of lateral YSV region of HH15 chick embryo after 15 minutes post-injection of FITC-Dex and 594-AcLDL. **(G, H)** Time-lapse images of FITC-Dex and 594-AcLDL in lateral YSV region. **(I)** Quantification of fluorescence intensity after FITC-Dex and 594-AcLDL injection. N=3 embryos were used for it. **(J, K)** Images of lateral YSV region after 15 minutes post-injection of Rho-LCA. (K) is magnified view. **(L)** Bright field images of lateral YSV region of HH15 chick embryo before and after Rho-LCA injection. Arrow heads indicate large erythrocyte aggregations. Scale bars: 500 μm in **B**, **C**, **J**, 20μm in **D**, **E**, **F**, **G**, **H**, **K**, **3**0μm in **K,** 1000μm in **L**.

In the quantitative evaluation of cancer cell extravasation, the visualization of blood vessels in fixed avian embryos has been a successful approach. However, while endothelial labeling in quail embryos is feasible using the QH1 antibody (Pardanaud et al., 1987), this method is not applicable in chick embryos due to their lack of reactivity to the QH1 antibody and the absence of suitable altaernative antibodies. To overcome this limitation, FITC-Dex, 594-AcLDL, or Rho-LCA were injected into the HH15-16 chick vasculature, followed by fixation to evaluate the preservation of vascular labeling. Although the fluorescence of FITC-Dex dissipated after fixation (data not shown), staining with 594-AcLDL and Rho-LCA remained intact (Fig. 3A, B). Post-fixation staining with Rho-LCA delineated a more continuous and distinct vascular wall compared to staining with 594-AcLDL (Fig. 3A, B). Therefore, for analyses in fixed chicken embryos, Rho-LCA staining appears to be more effective. Both Rho-LCA and 594-AcLDL are suitable for use in quail embryos as well (Fig. 3C, D); however, QH1 staining is recommended for labeling quail vascular endothelial labeling (Fig. 3E).

**Figure 3.**
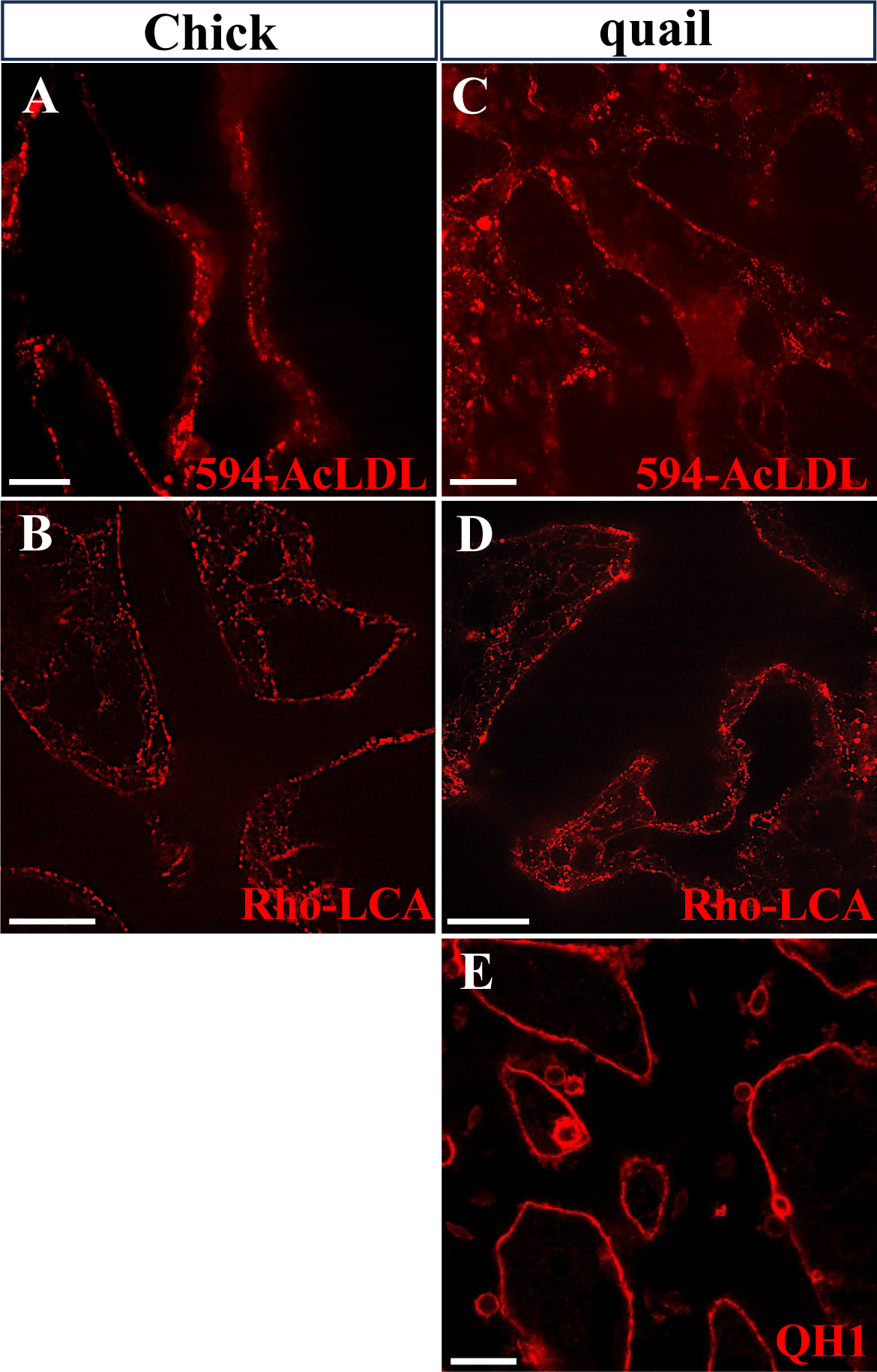
Vasculature visualization in fixed chick and quail embryo. **(A-E)** Images of lateral YSV regions of fixed HH15 chick and quail embryos stained by 594-AcLDL (A, C), Rho-LCA (B, D), and QH1 (E). Scale bars: 30 μm in **A-E.**

### Cancer cells extravasate in the YSV

To assess cancer cell extravasation from the YSV in chick and quail embryos at HH15, ZsGreen1-expressing HT-1080 cancer cells were introduced into the vasculature of both types of embryos, and their distribution pattern was examined after 6 hours. In chick embryos receiving HT-1080 cell transplantation, 53.33 ± 6.80% of the cells observed in the lateral region of the YSV were located outside the blood vessels (Fig. 4A, C). Similarly, in quail embryos undergoing the same procedure, 75.76 ± 4.73% of the cells were found outside the blood vessels (Fig. 4B, C). These findings suggest that the majority of HT-1080 cells undergo extravasation within 6 hours in the YSV of both avian species.

**Figure 4.**
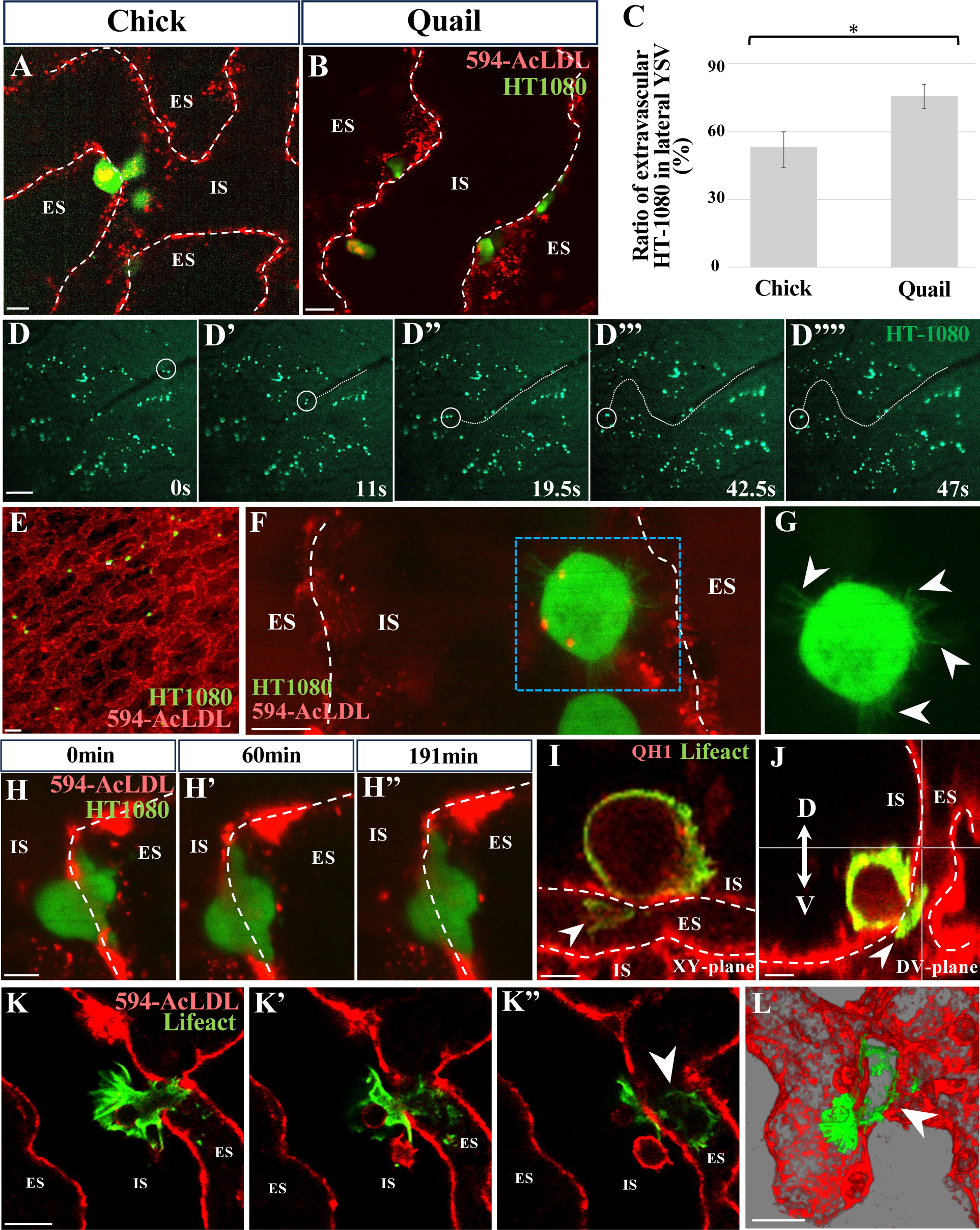
HT-1080 extravasate in the YSV. **(A, B)** Images of lateral YSV regions of HH15 chick (A) and HH15 Quail embryo (B) after 6hrs post-injection of ZsGreen1+ HT-1080 cells and 594-AcLDL. **(C)** Quantification of ZsGreen1+ HT-1080 cells present in the lateral YSV region of HH15 chick (N=3 embryos) and quail (N=3 embryos), specifically those located outside the blood vessels after 6hrs post-injection. *: P<0.05. **(D)** Time-lapse images of ZsGreen1+ HT-1080 cells in chick embryo after 1minutes post-injection. The white cercle and white dot line indicate the location of the cancer cells and their circulating trajectories, respectively. **(E)** Low magnification image of lateral YSV region after 1min post-injection of ZsGreen1+ HT-1080 cells and 594-AcLDL. **(F, G)** Magnified images of lateral YSV region after 30 minutes post-injection of ZsGreen1+ HT-1080 cells. (G) is magnified views of dotted box area shown in (F). White arrows indicate cellular protrusions. **(H)** Time-lapse images of an extravasating ZsGreen1+ HT-1080 cell in chick embryo after 1.5h post-injection. **(I, J)** Images of lateral YSV region of fixed HH15 quail embryo after 4hrs post-injection of Lifeact-expressing HT-1080 cells. White arrows indicate invadopodia-like protrusions. **(K, L)** Sequential confocal images of lateral YSV region (K) and their 3D reconstruction (L) of a fixed HH15 quail embryo after 6hrs post-injection Lifeact-expressing HT-1080 cells. White arrowheads indicate membrane bleb-like protrusions. White dotted lines indicate boundary between intravascular space (IS) and extravascular space (ES). Scale bars: 20μm in A, B, F, G, K, 300μm in D, 100μm in E, 10μm in H, I, J, L.

To capture the behavior of HT-1080 cells extravasating from the YSV region, we analyzed earlier time points after transplantation. Within 5 minutes post-transplantation, many HT-1080 cells ceased circulation at the lateral region of the YSV (Fig. 4D), indicating cancer cell arrest occurs. Most of these cells were observed adhering to the vessel wall (Fig. 4E-G), suggesting that adhesion is the main mechanism of HT-1080 arrest in the YSV. Interestingly, cancer cells were observed to form numerous microvillus-like cellular protrusions within intravascular space, facilitating their adhesion to the vessel wall (Fig. 4F, G).

At 1.5 hours post-transplantation, a significant portion of adherent HT-1080 cells on the vascular wall were observed initiating TEM (Fig. 4H). We identified two types of cellular protrusions employed by cancer cells during TEM: rod-like protrusions, reminiscent of invadopodia, which are supported by actin fibers (Fig. 4I, J), and balloon-like protrusions, resembling membrane blebs, with fewer actin fibers (Fig. 4K, L). These TEM events were primarily observed on the ventral side of the vascular wall in the YSV (Fig. 4J). Thus, the extravasation of cancer cells in the YSV was demonstrated, along with the capability to finely observe their cellular behaviors in both live and fixed embryos.

### Quantitative analysis of cancer cell extravasation in the YSV

The YSV presents an ideal setting for the quantitative assessment of cancer cell extravasation, owing to its small size and ease of observation. To gauge the efficacy of quantitative analysis within the YSV, we employed a functional inhibition model targeting Integrinβ1 (Itgnβ1), a molecule implicated in extravasation (Sokeland and Schumacher, 2019). Specifically, we induced the expression of a dominant-negative mutant of Itgnβ1, ΔC-Itgnβ1 (Lee et al., 2006), lacking the C-terminal domain, in HT-1080 cells. These ΔC-Itgnβ1+ HT-1080 cells, along with control mCheery+ cells, were uniformly transplanted into the vasculature of quail embryos (Fig. 5A). After 6 hours, the embryos were fixed, and the distribution of each cell population within the lateral region of the YSV vasculature was determined. In control experiments where HT-1080 cells expressing only ZsGreen1 or mCheery were transplanted, no discernible difference in extravasation rates between these cell populations was noted (Fig. 5B, C, F). However, our findings revealed a significantly reduced proportion of ΔC-Itgnβ1-expressing cancer cells located outside the YSV vasculature compared to controls (Fig. 5D, E, F). These results demonstrate the requirement for Itgnβ1 function in extravasation within the YSV and highlight the efficacy of the YSV analysis system in quantitatively assessing cancer cell extravasation.

**Figure 5.**
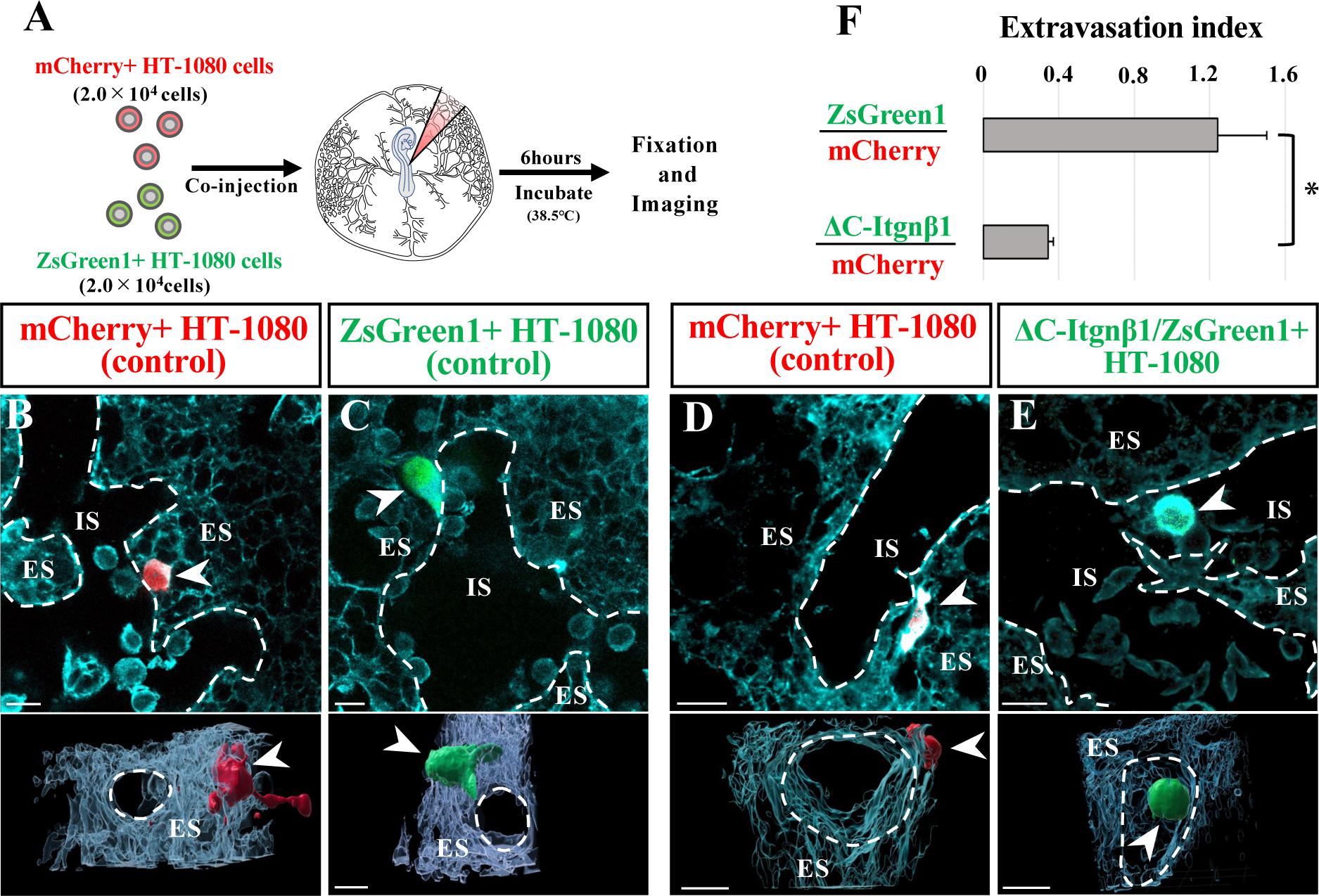
Quantitative analysis of cancer cell extravasation in the YSV. **(A)** Schematic representation of the experimental procedure of cancer cell transplantation. Each 20,000 mCherry+ and ZsGreen1+ HT-1080 cells are transplanted into HH15 quail embryo. **(B-E)** Confocal and 3D reconstruction images of lateral YSV regions of HH15 quail embryos after 6h post-injection. ECs are stained by QH1 (cyan). HT-1080 cells are labeled by mCherry (red) or ZsGreen1 (green). **(F)** Extravasation index of HT-1080 cells. The index is calculated as the ratio of extravascular mCherry+ cells among all mCheery+ cells in the lateral YSV, divided the ratio of extravascular ZsGreen1+ cells among all ZsGreen1+ cells. N=3 embryos were used for each group. *: P<0.05. Scale bars: 20μm in **B-E.**

## DISCUSSION

In this study, we explored the potential of avian YSV vasculature as a robust system for imaging and quantitatively analyzing extravasation. We anticipated several advantages with this analysis sytem in the following aspects: (1) Due to its planar nature, YSV forms a two-dimensional vascular network, facilitating easy live imaging at high resolution. (2) The developed *ex vivo* observation method further enhances high-resolution imaging capabilities (Kidokoro et al., 2018). (3) YSV is small in scale, allowing for comprehensive panoramic observation. Through our analysis, we established an effective YSV vascular labeling method for both live and fixed samples, characterized vascular structure and properties, and captured the temporal dynamic of HT-1080 behavior within the YSV. Leveraging the unique characteristics of the YSV, our whole-mount imaging analysis demonstrated the observation of fine structures, such as cancer cell protrusions, without the need for sectioning, enabling comprehensive visualization of intravital cancer cell behavior. The thinness of the structure also facilitated precise 3D reconstruction. Furthermore, utilizing this compact system, we found that quantitative measurement of cancer cell extravasation rates was remarkably straightforward.

### YSV as a cancer arrest model

We have demonstrated that the YSV possesses a relatively large minimum diameter of 15 μm and maintains integrity, preventing the passage of particles such as 250,000 MW Dextran. It exhibits a typical vascular structure comprising the layers of endothelium, BM, and SMC. Furthermore, the sparse distribution of the BM and SMC layer suggests classification closer to veins rather than capillaries. The arrest of HT-1080 cells in the YSV is primarily adhesive rather than occlusive, likely attributable to the larger diameter of the YSV compared to cancer cells. In contrast, the CAM and Zebrafish vascular models, where cancer cell arrest appears mainly occusive (Kim et al., 2016; Tobia et al., 2013), have smaller diameters than the YSV. Thus, it is possible that the YSVmodel could capture extravasation behaviors different from those observed in these existing models.

Cancer cell adhesion occurs within 5 minutes post-injection of HT-1080 cells into the vasculature, indicating a high affinity between the YSV and HT-1080 cells. Factors contributing to this affinity are presumed to be adhesion molecules or glycocalyx on the cell membrane (Offeddu et al., 2021; Reymond et al., 2013). Although the exact mechanism of adhesion remains unclear, it may involve fine cellular protrusion resembling microvilli. Future investigations focusing on functional analysis of adhesion molecules and molecules involved in microvilli formation are necessary to elucidate the cellular and molecular entities responsible for this affinity.

### YSV as a cancer TEM model

In our investigation, we observed TEM of HT-1080 cells within the YSV region. Cellular behaviors during TEM included invadopodia-like and bleb-like cellular protrusions. The invadopodia-like behavior resembled that observed in TEM across CAM (Leong et al., 2014), Zebrafish (Stoletov et al., 2007), and device-based systems (Wang et al., 2013). Conversely, while bleb-like behavior of cancer cells has been noted *in vitro* (Aoki et al., 2021; Halder et al., 2021), its occurrence during extravasation remains unreported. The YSV presents a promising platform for studying cancer cell bleb behavior during TEM, offering insights into how cancer cells distinguish between invadopodia and blebs.

TEM predominantly occurred on the ventral side of the vessel wall within the YSV, where the layers of basement membrane (BM) and smooth muscle cells (SMCs) were less developed. If the propensity for TEM and the vulnerability of the vascular structure are related, it raises significant questions about whether cancer cells have the ability to seek out locations conducive to TEM and, if so, what mechanisms underlie this ability. The YSV analysis system is expected to contribute to elucidating these questions.

Within 6 hours post-transplantation, the majority of HT-1080 cells complete TEM in YSV. In comparison, in CAM systems, approximately 30% undergo TEM within 6 hours post-transplantation (Leong et al., 2014), while in Zebrafish models, approximately 24hours are required for 30% to undergo extravasation (Chen et al., 2016a). Hence, the time for TEM to occur in the YSV system is notably rapid. The YSV analysis system not only serves as a specialized imaging tool but also offers an efficient analysis platform, saving both time and cost in therapeutic and pharmacological applications.

## ACKNOWLEDGEMENTS

For 3D image processing, we thank the Center for Advanced Instrumental and Educational Support of the Faculty of Agriculture, Kyushu University. This work was supported by the following grants: JSPS KAKENHI (Grant number 22H02634 for D. S.) and Princess Takamatsu Cancer Research Fund, Shinnihon Foundation of Advanced Medical Treatment Research, Terumo Life Science Foundation for D. S.

## AUTHOR CONTRIBUTIONS

M. Morita, R. F. and D. S. designed the study. M. Morita and R. F. performed most experiments and analyzed data. M. Morimoto and A. R. performed a part of the immunostaining experiments. T. T. performed 3D image acquisition. A. I., T. I., and J. I. prepare cancer cells. D. S., Y. H. and Y. A. supervised the study. D. S. wrote the manuscript.

